## Supplemental figures and methods for "Emergence of collective oscillations in adaptive cells"

Wang et al.

### Supplementary Figures

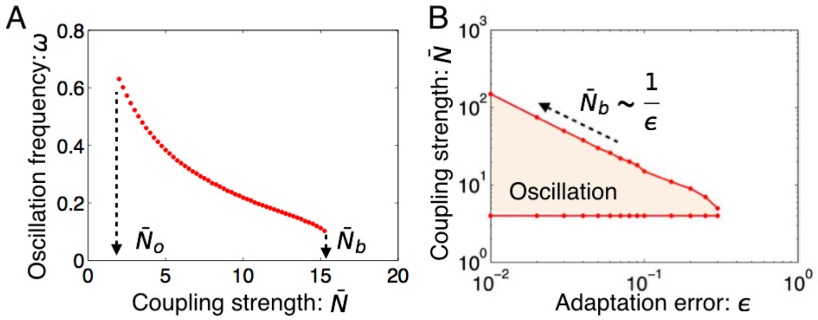


**Supplementary Figure 1. Collective oscillations in the model Equation (13) with nonlinear adaptation ($c_3 = 1$) and linear signal relaxation ($d_3 = 0$).** ($A$) Oscillation frequency against the effective cell density $\bar{N} = \alpha_2\alpha_1 N$. ($B$) The phase diagram in the plane spanned by $\bar{N}$ and the adaptation error $\epsilon$. Other parameters are the same as in Fig. 3 of the Main Text.

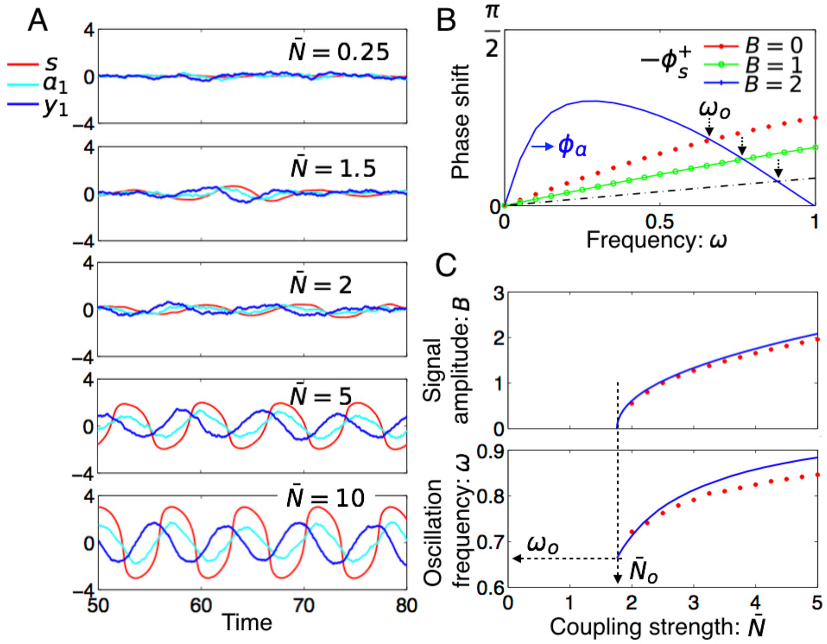


**Supplementary Figure 2. Collective oscillations in the model Equation (13) with linear adaptation ($c_3 = 0$) and nonlinear signal relaxation ($d_3 = 1$).** ($A$) Temporal trajectories at various values of $\bar{N}$. ($B$) The phase lead $\phi_a$ and lag $-\phi_s^+$ against $\omega$ at selected oscillation amplitudes. ($C$) The predicted oscillation frequency and amplitude as compared with those obtained from numerical simulations. Other parameters are the same as in Fig. 3 of the Main Text.

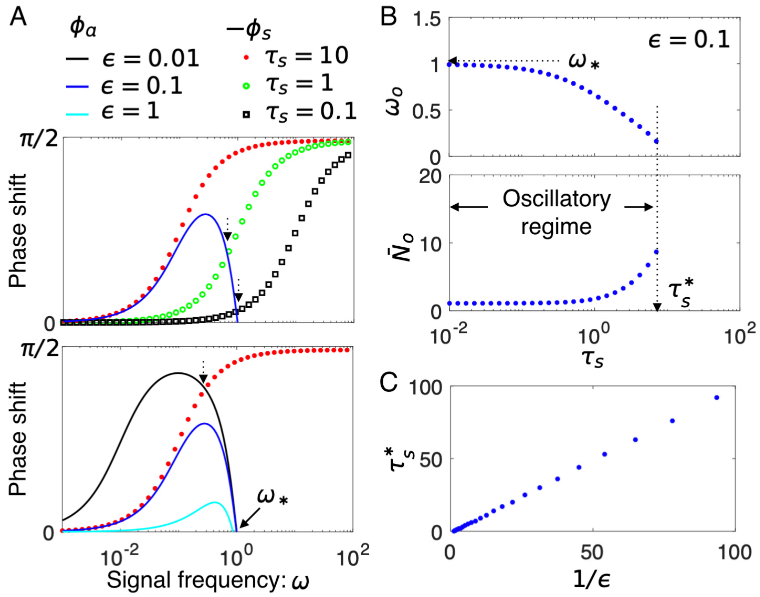


**Supplementary Figure 3**. Phase-matching at the onset of oscillations for different values of signal timescale $\tau_s$ and adaptation error $\epsilon$. (*A*) Phase shift of the cell activity ($\phi_a$) and signal response ($|\phi_s|$) at selected values of $\tau_s$ (upper panel) and adaptation error $\epsilon$'s (lower panel). The onset frequency is given by the intersection of the two curves [Equation (12a)]. (*B*) Predicted onset frequency $\omega_o$ and onset coupling strength $\bar{N}_o$ for different values of $\tau_s$. Oscillations will not be found when the signal relaxation time $\tau_s > \tau_s^*(\epsilon)$. (*C*) Numerical results supporting linear scaling between $\tau_s^*(\epsilon)$ and $1/\epsilon$. The data are obtained from the coupled adaptive circuits under the same parameters (except $\epsilon$ and $\tau_s$) as in Fig. 3 in the Main Text.

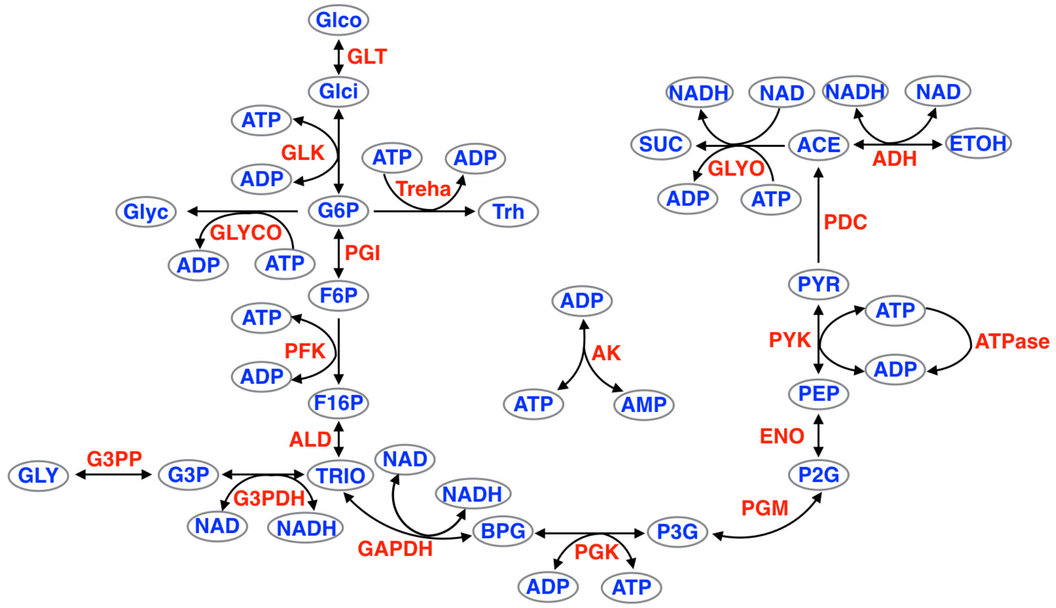


**Supplementary Figure 4**. **The network of reactions in a detailed model of glycolysis**[15]. Letters in blue denote metabolites, while those in red are the reactions. Directional (bidirectional) arrows indicate irreversible (reversible) reactions. Abbreviations: Glco, glucose; ACE, acetaldehyde, ADH, alcohol dehydrogenase; AK, adenylate kinase; ALD, fructose-1,6-bisphosphate aldolase; BPG, 1,3-bis-phosphoglycerate; ENO, phosphopyruvate hydratase; F16P, fructose-1,6-bisphosphate; F6P, fructose 6-phosphate; GAPDH, D-glyceraldehyde-3-phosphate dehydrogenase (phosphorylating); G3P, glycerol 3-phosphate; G3PDH, glycerol 3-phos- phate dehydrogenase; G6P, glucose 6-phosphate; GLYCO, glycogen branch; GLK, glucokinase (a hexokinase); P2G, 2-phosphoglycerate; P3G, 3-phosphoglycerate; PEP, phosphoenolpyruvate; PDC, pyruvate decarboxylase; PGI, glucose-6-phosphate isomerase; PFK, 6-phosphofructokinase; PGK, phosphoglycerate kinase; PGM, phosphoglycerate mutase; PYK, pyruvate kinase; PYR, pyruvate; Treha, trehalose branch; SUC, succinate branch; GLYO, glyoxylate shunt.

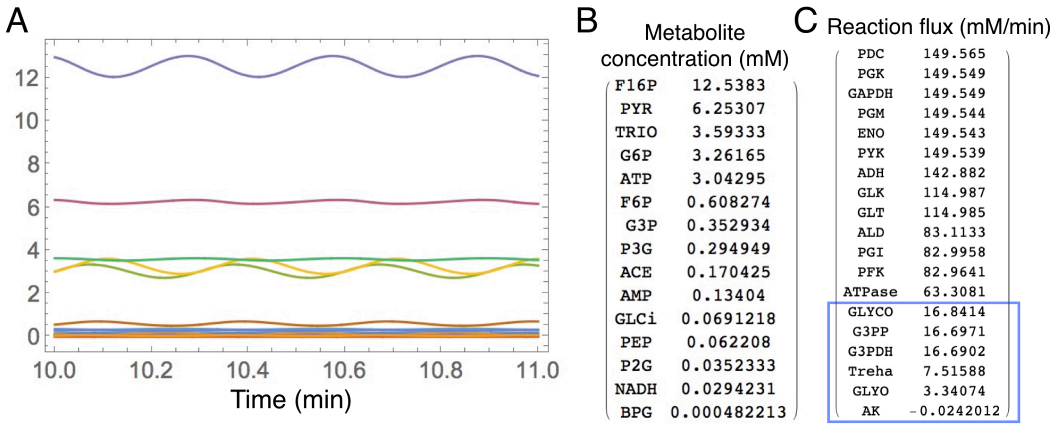


**Supplementary Figure 5**. **Spontaneous oscillations in the full model**. (*A*) Trajectories of all metabolites at glucose concentration Glco=10. (*B*) and (*C*) Time-averaged metabolite concentrations and reaction fluxes in descending order.

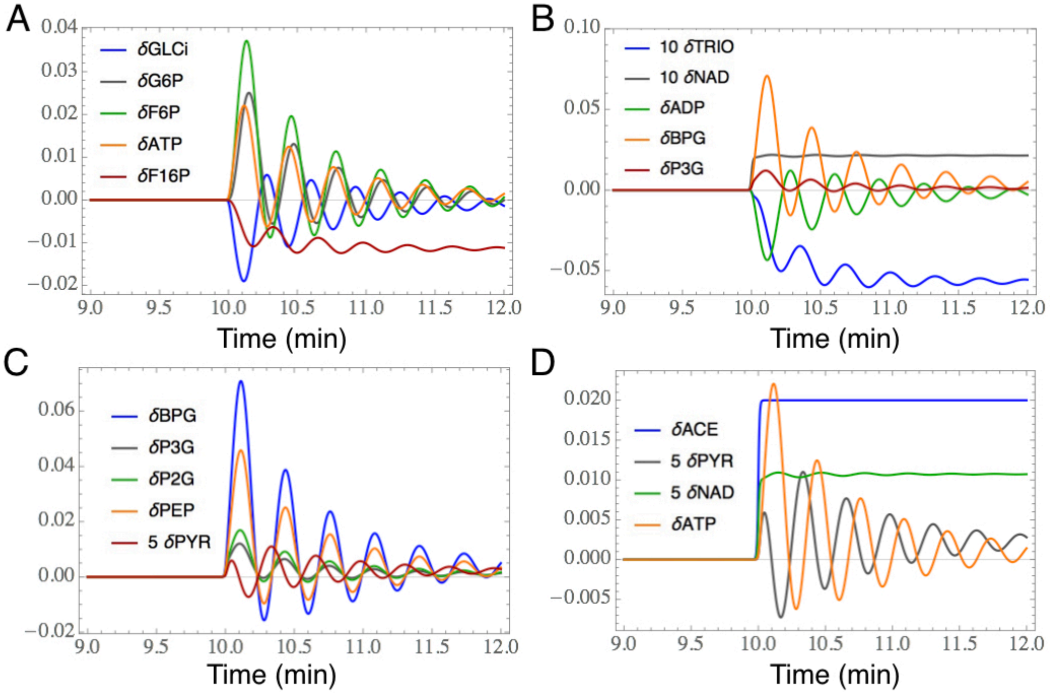


**Supplementary Figure 6**. **Response of metabolites to a redox signal at low ACE concentrations**. Here, $\text{ACE}(t) = \text{ACE}_0[1 + 0.02H(t)]$, with $H(t)$ being a Hill function with a large Hill coefficient. The notation $\delta x$ of a variable $x$ represents its relative change from a pre-stimulus level $\bar{x}$, i.e., $\delta x \equiv [x(t) - \bar{x}]/\bar{x}$. Quantities such as PYR and NAD which have too small values are amplified to make them visible on the plot. ($A$) The response of metabolites around G6P in the upper section of the glycolytic pathway; ($B$) The response of metabolites around BPG in the middle section of the glycolytic pathway; ($C$) The response of metabolites from BPG to PYR in the lower section of the glycolytic pathway; and ($D$) The response of metabolites in the downstream fermentation pathway. Parameters: $\text{ACE}_0 = 0.05$ and $\text{Glco} = 10$.

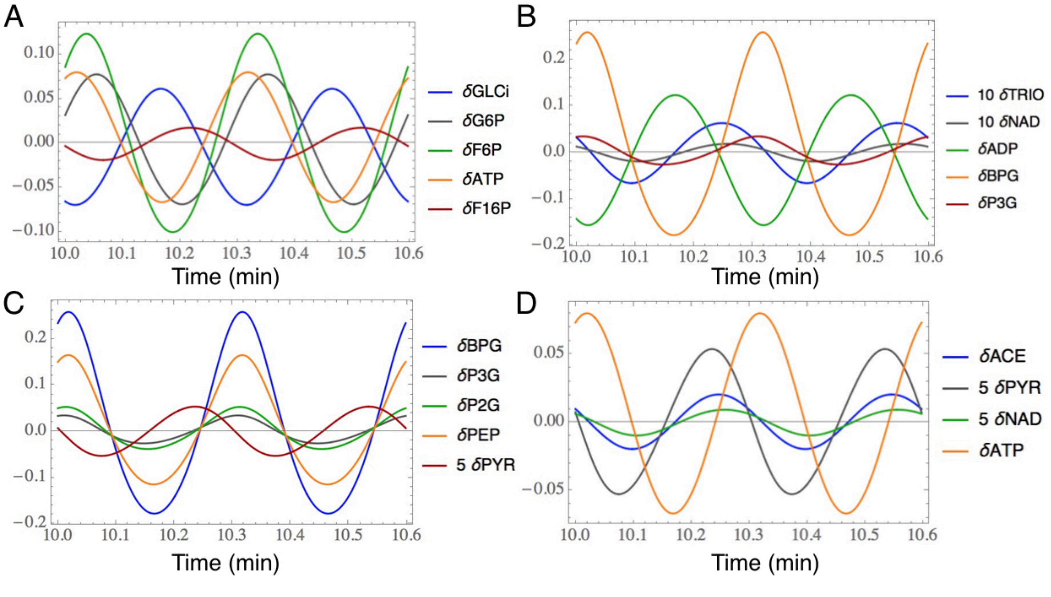


**Supplementary Figure 7**. **Concentration variations along the glycolytic pathway stimulated by a periodic redox signal**. Here, $\text{ACE}(t) = \text{ACE}_0[1+0.02\sin(\omega t)]$. Organization of metabolites in panels ($A$)-($D$) is the same as in Supplementary Figure **6**. Parameters: $\text{ACE}_0 = 0.05$, $\text{Glco} = 10$, and $\omega = 21$.

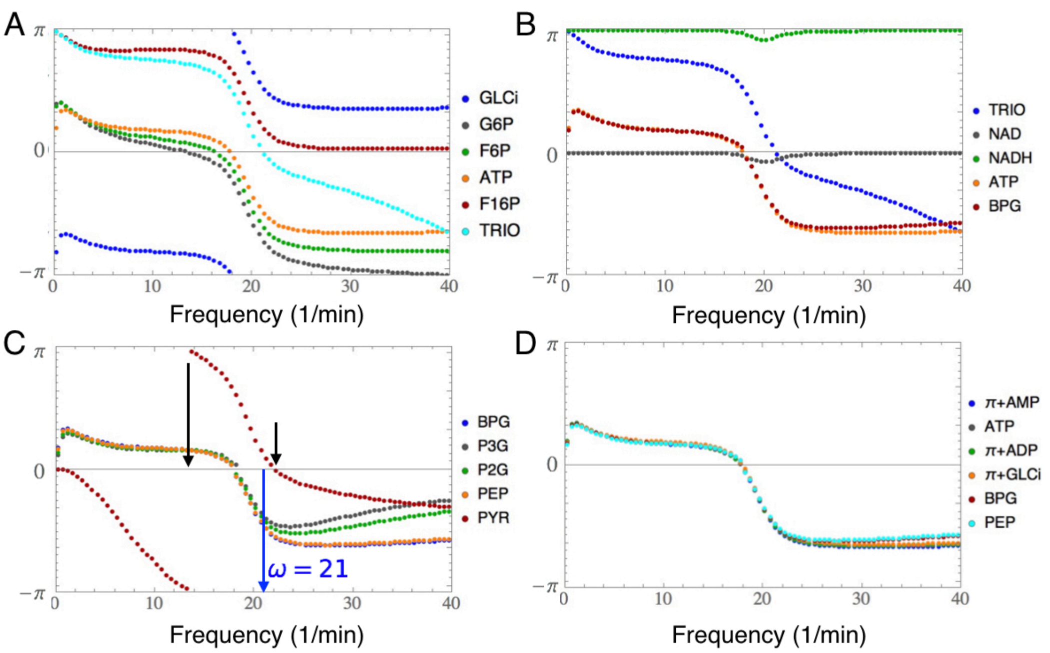


**Supplementary Figure 8. Phase shifts of metabolites against the frequency of a sinusoidal ACE signal**. (*A*) Metabolites in the "preparatory phase" of the glycolytic pathway, where ATP is consumed to activate the 6-carbon ring molecule. (*B*) Substrate, product and cofactors of the reaction GAPDH that act as the receptor of the redox signal, together with ATP. (*C*) Metabolites in the "payoff phase" of the glycolytic pathway, where ATP is harvested. For the particular values of the extracellular glucose and acetaldehyde chosen, phase lead of PYR over ACE occurs in the range of frequencies delimited by black arrows. The blue arrow indicates the intrinsic frequency studied in Supplementary Figure **7**. (*D*) Metabolites that appear in Equation (25). The phase shift of a number of metabolites shows a dip at the low frequency end, indicating a small but finite adaptation error. Parameters: $\mathrm{ACE}_0 = 0.05$, $\mathrm{Glco} = 10$.

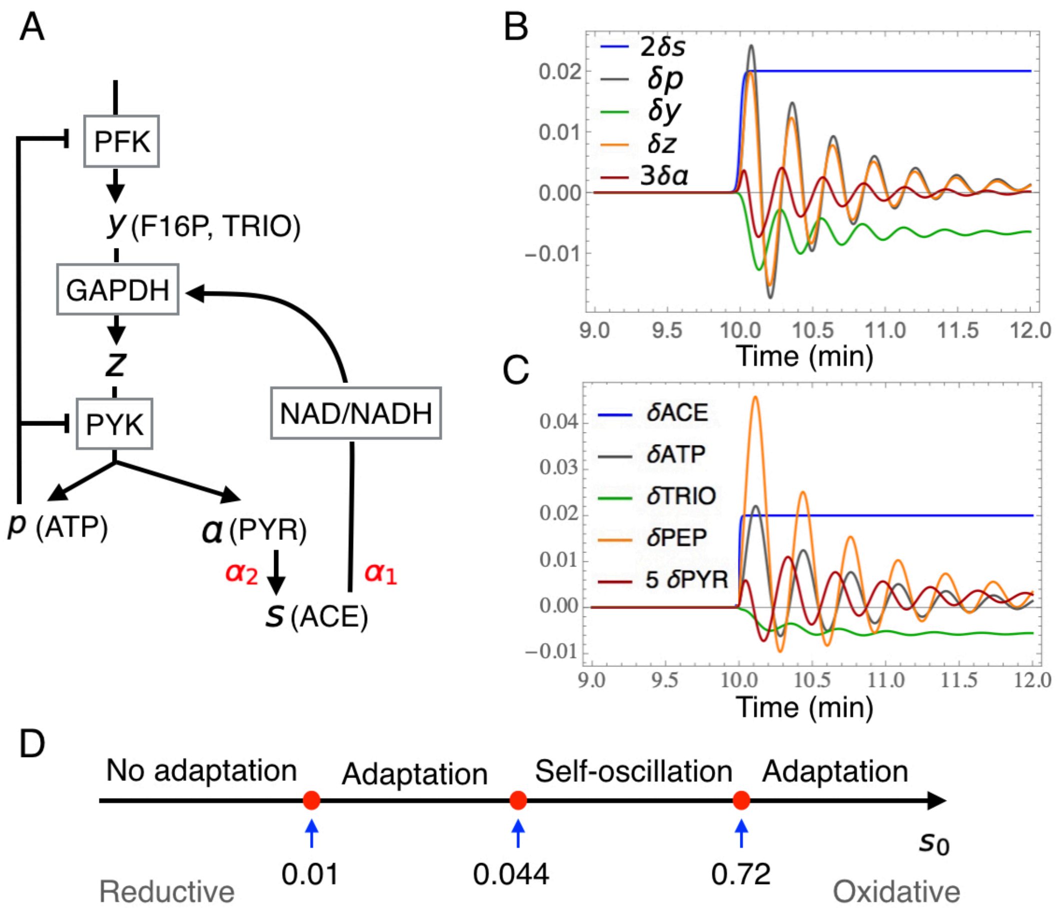


**Supplementary Figure 9**. **A minimal model of glycolysis with redox control**. (*A*) The network of metabolites (symbols) and reactions (boxes). (*B*) Response of metabolites upon a stepwise perturbation $s(t) = 0.04(1 + 0.01H(t))$. (*C*) Response of corresponding metabolites in the full model computed using parameter values given in Supplementary Figure **6**. (*D*) Response properties of the minimal model as the intracellular redox state changes from reductive to oxidative (left to right). Parameters: $h = 3, \alpha_2 = \alpha_1 = 1$, $\tau = 0.01$, $c_0 = 0.02$, and $\epsilon = 0.01$.

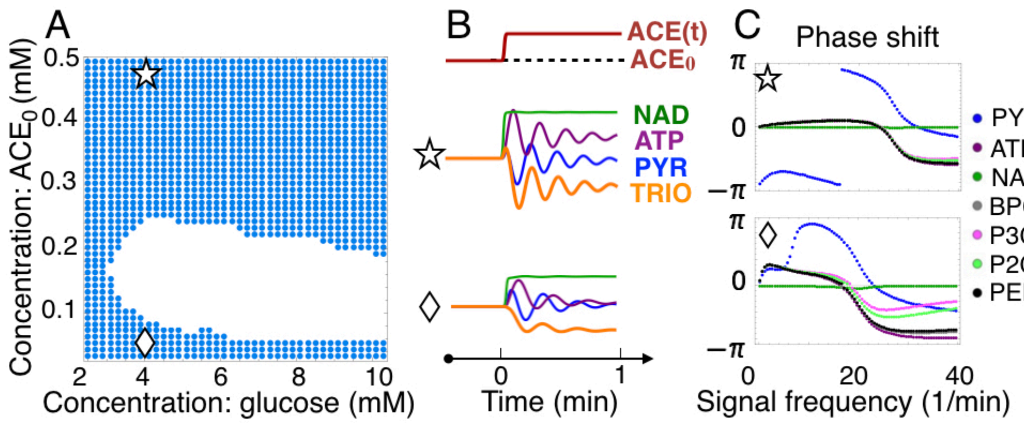


**Supplementary Figure 10**. **A mutant glycolysis model with the GLYO reaction switched off**. (*A*) Phase diagram of this model at an environment set by the glucose and ACE concentration. The white regime is the self-oscillatory regime, while the blue one is the adaptive regime for both ATP and PYR. The adaptive regime of PYR and ATP overlaps now, in agreement with the minimal model. (*B*) The temporal dynamics of selected metabolites under a shift of the signal ACE at the two indicated conditions (star and diamond in *A*). (*C*) The phase shift of various metabolites under a periodic perturbation of the signal ACE at the selected conditions.

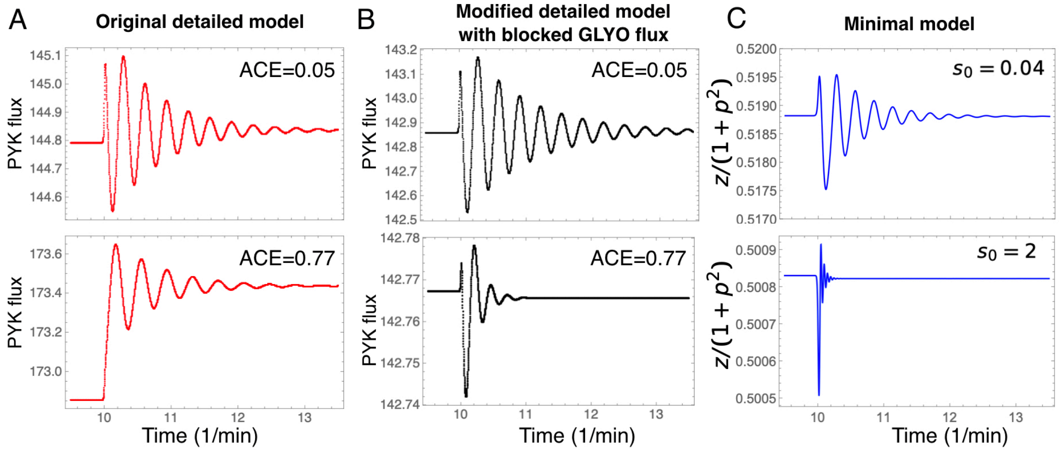


**Supplementary Figure 11**. **The response of PYK flux to a step perturbation of ACE at** $t = 10$. ($A$) The response of the original full model at ACE=0.05 and 0.77, respectively. ($B$) The response of the modified full model with blocked GLYO reaction at ACE=0.05 and 0.77, respectively. ($C$) The response of the minimal model. Parameters: Glco=10 for ($A$) and ($B$); the parameters for the minimal model are the same as in Supplementary Figure **9**.

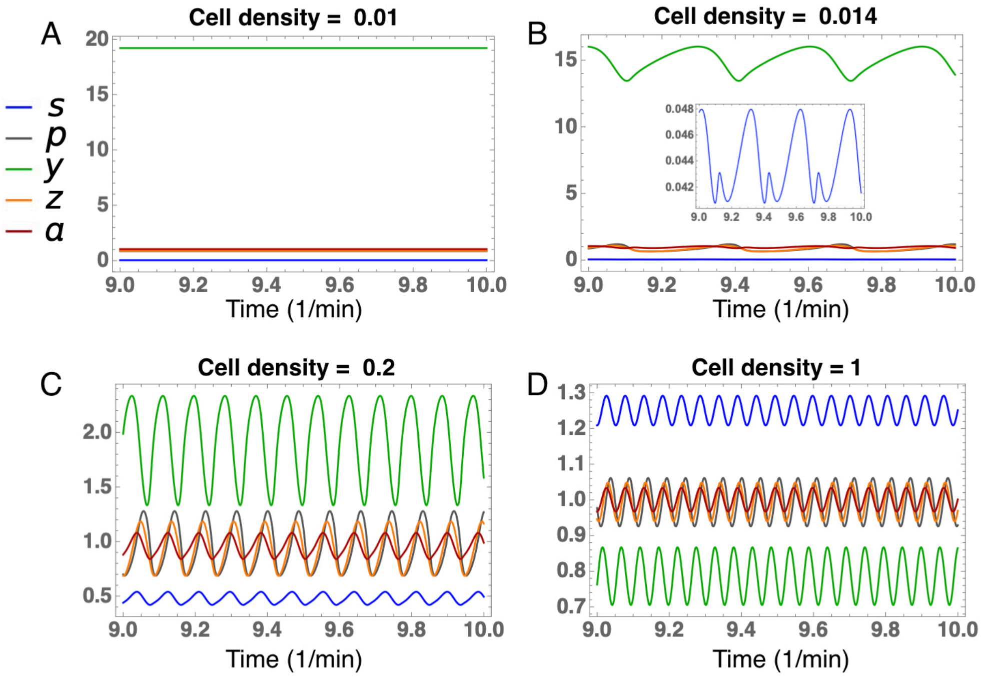


**Supplementary Figure 12**. **Collective dynamics of the minimal model coupled via Equation (28)**. ($A$)-($D$) Temporal trajectories at selected values of the cell density $\rho$. The same color scheme of variables is used. Inset in $B$ shows the signal trajectory on an enlarged scale. Parameters: $h = 3, \alpha_2 = \alpha_1 = 1, \epsilon = 0.01$, $k_{\rm in} = 0.5$, $k_{\rm ex} = 0.3$, $\tau = 0.01$, $c_0 = 0.02$, and $\tau_s = 0.001$.

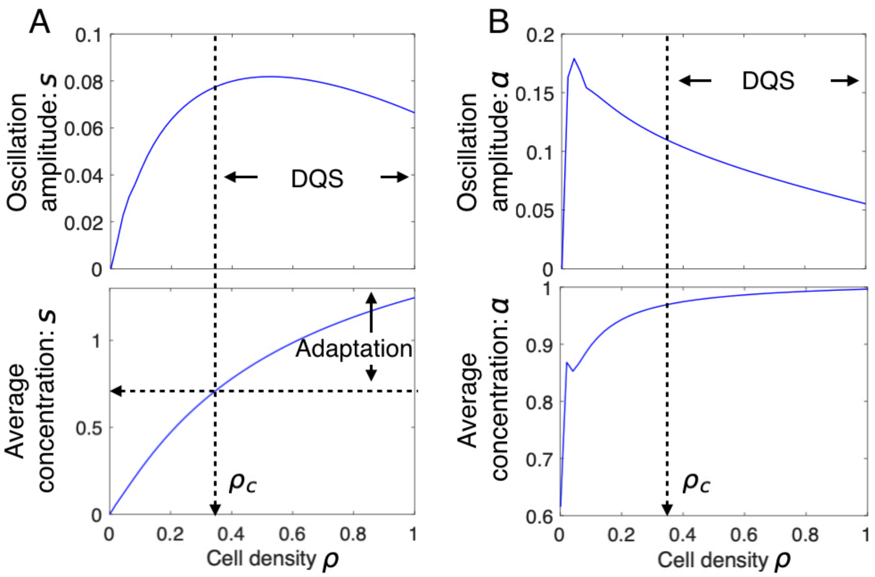


**Supplementary Figure 13**. **Collective oscillations against the yeast cell density**. (*A*) Upper panel: oscillation amplitude of the signal $s$ as a function of the cell density $\rho$. Lower panel: time-averaged signal concentration against $\rho$. At $\rho_c = 0.34$, the signal strength reaches the upper threshold $s_c = 0.72$ for the oscillating state of individual cells (Supplementary Figure **9***E*). The oscillating state at $\rho > \rho_c$ can be considered as DQS driven by adaptation. (*B*) Upper panel: oscillation amplitude of the sender node $a$ against $\rho$. Lower panel: time-averaged value of $a$ against $\rho$. Parameters are the same as in Supplementary Figure **12**.

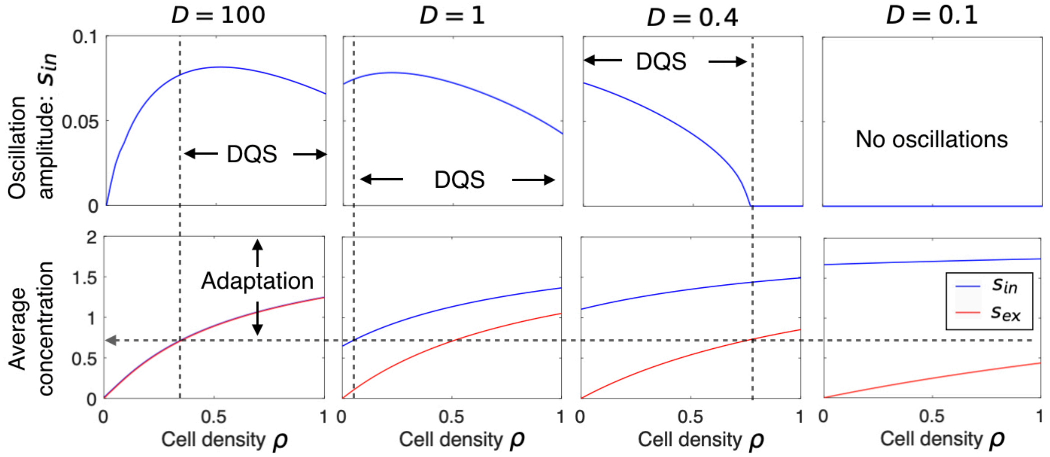


**Supplementary Figure 14**. **Effect of delay in cross-membrane transport of the signalling molecule on collective dynamics**. Results of numerical integration of Eqs. (26)] coupled to the two-component signal dynamics Equation (27)] at selected values of $D = 100, 1, 0.4, 0.1$. In the lower panels, the blue and orange dots correspond to the average of $s_{\mathrm{in}}$ and $s_{\mathrm{ex}}$, respectively. Other parameters are the same as in Supplementary Figure **12**.

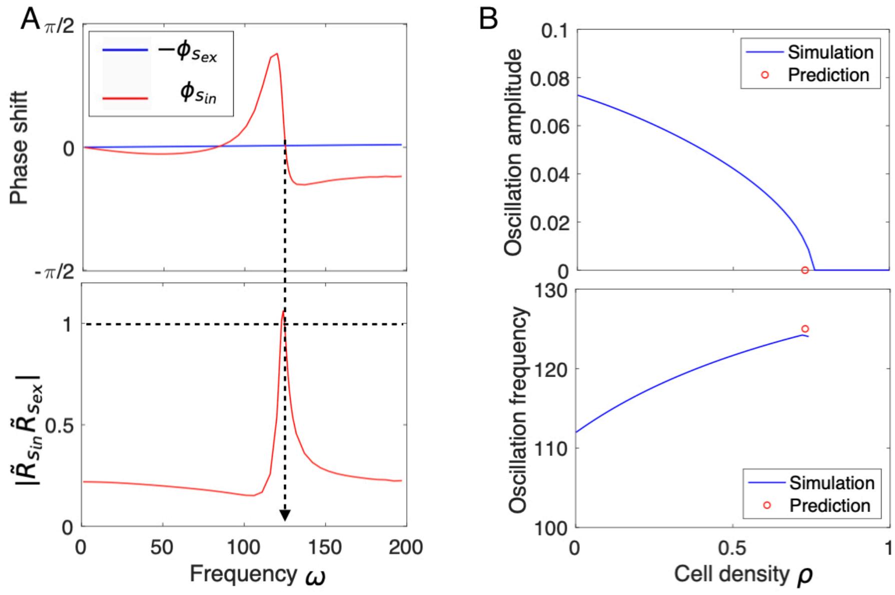


**Supplementary Figure 15**. **Predicting the death of collective oscillations at large cell densities**. ($A$) Predicting the critical frequency using self-consistency check. Upper panel: the spectrum of the numerically computed phase shift of the intracellular signal $s_{\text{in}}$ and the extracellular signal $s_{\text{ex}}$, respectively. Lower panel: the spectrum of the compound gain $|\tilde{R}_{s_{\text{in}}}\tilde{R}_{s_{\text{ex}}}|$. Both of the spectra are computed at the upper critical cell density $\rho_c = 0.73$ and $D = 0.4$. ($B$) Comparing the prediction with results from direct numerical simulation of population dynamics [Eqs. (26) and Equation (27)] at $D = 0.4$. Other parameters are the same as in Supplementary Figure **12**.

**Supplementary Note 1. Nonequilibrium thermodynamics of adaptive response**

*Phase-leading response of an adaptive variable*

Consider the temporal variation $a_t$ of an intracellular observable $a$ induced by a sinusoidal signal $s_t$ from the environment at frequency $\omega$. The amplitude of $s_t$ is assumed to be small, so that it can be treated as a perturbation. The observable $a$ is either directly or indirectly coupled to the signal $s$. On the scale of a single cell, both $a_t$ and $s_t$ may contain stochastic components. In the following, we shall examine the noise averaged response of $a$ to the deterministic part of $s$, i.e., the signal. As usual, we use $\langle\cdot\rangle$ to denote the noise average. The following convention on forward and inverse Fourier transforms is adopted,

$$\tilde{f}(\omega) = \int_{-\infty}^{\infty} f(t)\exp(i\omega t)dt, \quad f(t) = \int_{-\infty}^{\infty} \tilde{f}(\omega)\exp(-i\omega t)\frac{d\omega}{2\pi}. \tag{1}$$

In a steady-state, the ratio of the Fourier amplitudes $\langle\tilde{a}(\omega)\rangle$ and $\langle\tilde{s}(\omega)\rangle$ defines the response function $\tilde{R}_a(\omega) \equiv \langle\tilde{a}(\omega)\rangle/\langle\tilde{s}(\omega)\rangle$, which can be separated into its real $\tilde{R}'_a$ and imaginary $\tilde{R}''_a$ parts. (For a cellular variable $a$ that follows stochastic dynamics, the ensemble averaged response is considered.) The well-known Kramers-Krönig relation from causality requirement on the response function states[2]:

$$\tilde{R}'_a(\omega) = \frac{2}{\pi}\int_0^{\infty} \tilde{R}''_a(\omega_1)\frac{\omega_1}{\omega_1^2-\omega^2}d\omega_1. \tag{2}$$

In the case of a perfectly adapting $a$, the response vanishes under a sufficiently slow stimulus, i.e., $\lim_{\omega\to 0}\tilde{R}_a(\omega) = 0$. Equation (2) then requires,

$$\int_0^{\infty} \tilde{R}''_a(\omega_1)\omega_1^{-1}d\omega_1 = 0. \tag{3}$$

Consequently, $\tilde{R}''_a(\omega)$ must change sign at least once along the frequency axis. Let $\phi_a = -\arg(\tilde{R}_a)$ be the phase of $-\tilde{R}_a(\omega)$, with the minus sign introduced by convention. Positive and negative values of $\tilde{R}''_a$ thus translate to phase-lag ($-\pi < \phi_a < 0$) and phase-lead ($0 < \phi_a < \pi$) of $a_t$ over $s_t$, respectively. By virtue of continuity, the sign change of $\tilde{R}''_a(\omega)$ is also expected in the partially adaptive case, provided the adaptation error $\epsilon \simeq \tilde{R}'_a(0)$ is sufficiently small.

*Energy outflow from an adaptive channel*

Auto-induced collective oscillations in a dissipative medium require an energy source. Below, we show that an active cell is able to output energy to a fluctuating $s$ in the presence of an adaptive channel. The power of the output depends on the strength of the coupling as well as the amplitude and frequency of the fluctuating signal.

Consider a slightly more general form of Equation (1) in the Main Text where the contribution from cell $j$ to the total thermodynamic force on $s$ is given by $O(a_j)$, which in general is nonlinear in $a_j$. The work done on $s$ by the cell in a time interval $(0, L)$ can then be written as

$$W_j = \int_0^L O_t\dot{s}_t dt, \tag{4}$$

where $O_t \equiv O(a_j(t))$ and $s_t$ are both fluctuating quantities in general. We now consider a sinusoidal signal $s_t = s_0 + \Delta s\cos(\omega t)$ with a small amplitude $\Delta s$. To the first order in $\Delta s$, we have

$$O_t \simeq O_t^{(0)} + \Delta s|\tilde{R}_O(\omega)|\cos\big(\omega t + \phi_O(\omega)\big). \tag{5}$$

Here $O_t^{(0)}$ denotes the stochastic trajectory of $O$ in the absence of the sinusoidal signal. As usual, the linear response function $R_O$ in the steady-state (ss) to a weak time-varying signal $s_t$ is introduced through

$$\langle O_t\rangle \simeq \langle O\rangle_{ss} + \int_{-\infty}^{t} R_O(t-\tau)s_\tau d\tau, \tag{6}$$

where $\langle\cdot\rangle$ denotes average over noise. The phase angle $\phi_O(\omega) \equiv -\arg \tilde{R}_O(\omega)$. Substituting expression (5) into Eq. (4) and taking the limit $L \to \infty$, we obtain the time-averaged output power from the cell through this channel (omitting the subscript $j$),

$$\overline{\dot{W}} = \overline{O_t \dot{s}_t} + o\big((\Delta s)^2\big) = \frac{1}{2}\omega|\tilde{R}_O(\omega)| \sin\phi_O(\omega)(\Delta s)^2 + o\big((\Delta s)^2\big). \tag{7}$$

Here the overline bar indicates averaging over time, and $o\big((\Delta s)^2\big)$ denotes terms higher than second order in $\Delta s$.

Given the relation $O_t = O(a_t)$, adaptation of the cellular variable $a$ to a slow-varying $s$ also implies the adaption of $O$ to the signal. The causality condition (2) applied to $O_t$ then requires $\tilde{R}''_O(\omega) \equiv -|\tilde{R}_O(\omega)| \sin\phi_O(\omega) < 0$ in a certain frequency range. Consequently, Eq. (7) predicts energy outflow from the channel under a periodic stimulation at these frequencies.

The discussion leading to Eq. (7) in the previous section can be easily extended to the energy outflow under an arbitrary signal variation $s_t$ with a power spectrum $\tilde{C}_s(\omega)$,

$$\overline{\dot{W}} = -\int d\omega\omega \tilde{R}''_O(\omega)\tilde{C}_s(\omega) + o(\Delta s^2), \tag{8}$$

where $\Delta s$ sets the overall amplitude of signal variation. If the cell were in thermal equilibrium, $a$ would respond passively to a time-varying signal with a phase-lag and dissipate the energy inflow generated by the stimulation. An adaptive cell, on the other hand, is able to output energy in the form of work when stimulated in the right frequency range. This form of energy outflow is different from the heat dissipation arising from keeping the system out of equilibrium as studied in Refs.[3–5].

*The Fluctuation-Dissipation Theorem*

The fluctuation-dissipation theorem (FDT) is generally presented as an identity between the response function of a chosen variable to an external perturbation and the correlation function of the variable in question with the one that is conjugate to the perturbation[6]. For Markov systems which are of interest here, FDT holds when the detailed balance condition on the state-space transition rates is fulfilled. We refer the reader to Refs.[7,8] for a detailed discussion, including more rigorous definitions of various quantities of interest.

Assuming that the signal $s$ affects the cell through coupling to a conjugate variable $O$ which is proportional to the variable $a$ of interest, i.e., $O = c_0 a$ with $c_0$ a proportionality constant. In this case, FDT states that

$$\tilde{R}''_O(\omega) = \frac{\omega\tilde{C}_O(\omega)}{2T} > 0. \tag{9}$$

Here, $\tilde{C}_O(\omega) = c_0^2\langle|\tilde{a}(\omega)|^2\rangle$ is the power spectrum of $O_t$ which is always positive. Equation (9) contradicts (3), re-affirming that receptor adaptation cannot be realised without the presence of active processes inside the cell.

In Ref.[9], adaptation through a 3-node incoherent feed-forward motif was considered. It was later shown that the topology even supports adaptation in an equilibrium setting[10]. The main difference between these models and the adaptive receptor model in the Main Text (Fig. 3 and Methods) is that, in the former, $s$ not only couples to $a$ directly, but also to other intracellular variables. The conjugate variable $O$ is then a combination of $a$ and other intracellular variables. We leave a detailed investigation of this issue to future work.

### Supplementary Note 2. A self-consistent scheme for frequency selection and oscillation amplitude determination

The thermodynamic analysis in the preceding section suggests the possibility of a positive feedback loop formed by a periodic signal and adaptive cells under generic conditions. Collective oscillations emerge when signal amplification by active cells overtakes signal dissipation in the passive medium. In this section, we examine this process in further detail and derive equations that can be used to determine the frequency and amplitude of auto-induced oscillations when the instability takes place. For simplicity, we shall consider a situation where diffusion of the signalling molecules in the medium is very fast so that spatial variations of $s$ is suppressed. Consequently, the notion of a well-defined transition to the oscillating state can be introduced.

*The phase matching condition and threshold cell density*

Given that individual cells couple to each other only through the signal field $s$, a self-consistency procedure similar to the solution of mean-field models in statistical physics can be employed. In this case, the linear equations governing an eigenmode with eigenvalue $\lambda$ can be divided into subgroups associated with individual cells. The internal variables of a given cell appear in one and only one of the subgroups. Solution of the subset of equations for cell $j$ yields the cell activity $\langle \tilde{a}_j \rangle = \tilde{R}_{a,j}(i\lambda)\langle \tilde{s} \rangle$. The function $\tilde{R}_{a,j}(i\lambda)$ is the same function introduced in the preceding section to describe the linear response of $a_j$ to a sinusoidal perturbation at frequency $\omega = i\lambda$. Likewise, a response function $\tilde{R}_s(\omega)$ from the linearised relaxational dynamics of $s$ can be obtained, treating contributions from cells as source terms, as in Eq. (1) of the Main Text. Combining the two steps, we arrive at the following eigenvalue equation for $\lambda$,

$$\tilde{R}_s(i\lambda) \sum_{j=1}^{N} \alpha_1 \tilde{R}_{a,j}(i\lambda) = 1. \tag{10}$$

When a particular eigenvalue crosses the imaginary axis, its real part vanishes, while its imaginary part $\omega_o$ (the onset frequency) satisfies,

$$\alpha_1 N_o \tilde{R}_s(\omega_o) \tilde{R}_{\bar{a}}(\omega_o) = 1. \tag{11}$$

Here $\tilde{R}_{\bar{a}}(\omega) \equiv N^{-1} \sum_{j=1}^{N} \tilde{R}_{a,j}(\omega)$ is the averaged single-cell response function.

Equation (11) can be written separately for the phase shift $\phi = -\arg \tilde{R}$ and amplitude $|\tilde{R}|$ of the response functions. For $\alpha_1 > 0$, we have,

$$\phi_{\bar{a}}(\omega_o) = -\phi_s(\omega_o), \tag{12a}$$

$$N_o = \frac{1}{|\alpha_1 \tilde{R}_s(\omega_o)||\tilde{R}_{\bar{a}}(\omega_o)|}. \tag{12b}$$

Eq. (12a) determines the frequency $\omega_o$ at the onset of collective oscillations, while Eq. (12b) gives the threshold cell density $N_o$. As we mentioned in the Main Text, when the signal is passive, phase lead by the cell is required for Eq. (12a) to be fulfilled. The explicit relation presented here complements the energy argument based on Eq. (7), with the activity-generated thermodynamic force $O_t$ being proportional to $\alpha_1 a_j$.

As it stands, the cell density $N$ does not appear explicitly in the phase-matching condition (12a). Therefore the frequency of collective oscillations can be estimated from separate measurements of the single-cell response and the medium response. In reality, it is conceivable that properties of the medium are affected by the presence of cells, e.g., the concentration of the signalling molecules secreted. Consequently, both $\tilde{R}_s(\omega)$ and $\tilde{R}_{\bar{a}}(\omega)$ may have certain weak dependence on $N$.

*The amplitude equations and frequency shift*

Beyond the initial instability, nonlinear effects need to be treated explicitly to determine the amplitude and frequency of oscillations. Assuming a periodic state, the signal strength $s(t)$ can be expressed as a Fourier series that includes the first harmonic as well as higher order harmonics produced by nonlinearities in the system dynamics. Likewise, the noise-averaged cellular activity $\langle a_j(t) \rangle$ can also be expressed as a Fourier series in $t$ with the same basic frequency. For weak noise, the trajectory of the system falls on a well-defined limit cycle whose mean radius $r$ sets the overall amplitude of oscillations, while the amplitude of the $n$th order harmonic scales as $r^n$. This structure allows for a systematic determination of the amplitudes using perturbation theory. Below, we illustrate the procedure in the case of cubic nonlinearities in both the dynamics for $s$ and the dynamics for $a$, and comment on similarities and differences in more general situations. When the cell's activity is noisy, more sophisticated schemes based on the probability distribution function of the cellular state need to be introduced (see, e.g. Ref.[11]).

Let us consider a noiseless version of the adaptive dynamics defined in the Main Text (Fig. 3 and Methods), together with a modified version of Eq. (1) that includes a cubic nonlinearity,

$$\tau_a \dot{a}_j = -(a_j - y_j) - c_3 a_j^3 + \alpha_2 s \tag{13a}$$

$$\tau_y \dot{y}_j = -(a_j + \epsilon y_j) \tag{13b}$$

$$\tau_s \dot{s} = -s - d_3 s^3 + \alpha_1 \sum_{j=1}^{N} a_j. \tag{13c}$$

Here $\tau_s = \gamma_s/K_s$ gives the relaxation timescale for the signal. We also set $\alpha_1 \to K_s\alpha_1$ for notational simplicity. The two coefficients $c_3$ and $d_3$ set the strengths of nonlinearities in the cellular and signal dynamics, respectively. The model has the inversion symmetry $s \to -s$ and $(a_j, y_j) \to (-a_j, -y_j)$, all $j$. Furthermore, if we redefine the sign of $s$ and at the same time change the sign of $\alpha_2$ and $\alpha_1$, the equations remain the same.

We now seek a periodic solution to Eqs. (13) in Fourier form,

$$s(t) = B\cos(\omega t) + \sum_{n=2}^{\infty} B^{(n)} \cos(n\omega t + \phi_s^{(n)}), \tag{14a}$$

$$a_j(t) = A_j \cos(\omega t + \phi_{a,j}) + \sum_{n=2}^{\infty} A_j^{(n)} \cos(n\omega t + \phi_{a,j}^{(n)}), \qquad j = 1, \dots, N, \tag{14b}$$

$$y_j(t) = C_j \cos(\omega t + \phi_{y,j}) + \sum_{n=2}^{\infty} C_j^{(n)} \cos(n\omega t + \phi_{y,j}^{(n)}), \qquad j = 1, \dots, N. \tag{14c}$$

The amplitudes and phase shifts, all assumed to be real, satisfy a set of equations which can be derived by substituting Eqs. (14) into Eqs. (13), and grouping terms according to the order of the harmonic.

Starting from the first harmonic in the expressions (14), the cubic terms in Eqs. (13a) and (13c) generate the first and third order harmonics according to the identity $(\cos\phi)^3 = (3\cos\phi + \cos 3\phi)/4$. Hence terms such as $A_j^3$ and $B^3$ are present in the equations for the first harmonic. On the other hand, the cubic nonlinearities do not generate even order harmonics if they are not included in the series initially. Hence, up to the third order in the amplitudes, the equations for the coefficients of the first harmonic take the form,

$$-i\omega\tau_a\tilde{a}_j \simeq -(1 + \frac{3}{4}c_3|\tilde{a}_j|^2)\tilde{a}_j + \tilde{y}_j + \alpha_2\tilde{s}, \tag{15a}$$

$$-i\omega\tau_y\tilde{y}_j = -\tilde{a}_j - \epsilon\tilde{y}_j, \tag{15b}$$

$$-i\omega\tau_s\tilde{s} \simeq -(1 + \frac{3}{4}d_3|\tilde{s}|^2)\tilde{s} + \alpha_1 \sum_j \tilde{a}_j. \tag{15c}$$

Here we have introduced the short-hand notations $\tilde{s} = B, \tilde{a}_j = A_j \exp(-i\phi_{a,j})$, and $\tilde{y}_j = C_j \exp(-i\phi_{y,j})$.

To gain an intuitive understanding of the oscillatory solution as the cell density increases beyond the threshold $N_o$, we first eliminate the intracellular variable $\tilde{y}_j$ in Eqs. (15a) and (15b) to obtain,

$$\tilde{a}_j = \tilde{R}_{a,j}^{+}(\omega)\tilde{s}, \tag{16}$$

where

$$\tilde{R}_{a,j}^{+}(\omega) \equiv \frac{\tilde{a}_j(\omega)}{\tilde{s}(\omega)} \simeq \frac{\alpha_2}{1 + 3c_3|\tilde{a}_j|^2/4 - i\omega\tau_a - 1/(i\omega\tau_y - \epsilon)} \tag{17}$$

is a “nonlinear response function” which expresses the ratio of the complex amplitudes of the first harmonic on the limit cycle. Similarly, Eq. (15c) can be rewritten as

$$\tilde{s} = \tilde{R}_s^{+}(\omega) \sum_{j=1}^{N} \alpha_1\tilde{a}_j, \tag{18}$$

where

$$\tilde{R}_s^{+}(\omega) \simeq \frac{1}{1 + 3d_3|\tilde{s}|^2/4 - i\omega\tau_s} \tag{19}$$

is a “nonlinear response function” of $s$ on the limit cycle. It is easy to see that $\tilde{R}_a^{+}(\omega)$ and $\tilde{R}_s^{+}(\omega)$ reduce to their respective linear counterparts $\tilde{R}_{a,j}(\omega)$ and $\tilde{R}_s(\omega)$ when the oscillation amplitudes vanish.

We now combine Eqs. (16) and (18) to obtain the self-consistency condition,

$$\alpha_1 N \tilde{R}_s^{+}(\omega)\tilde{R}_{\bar{a}}^{+}(\omega) = 1, \tag{20}$$

which is reminiscent of Eq. (11). Here $\tilde{R}_{\bar{a}}^{+}(\omega) \equiv N^{-1}\sum_{j=1}^{N} \tilde{R}_{a,j}^{+}(\omega)$ is the averaged single-cell nonlinear response function. When all cells are identical, $\tilde{R}_{\bar{a}}^{+}(\omega) = \tilde{R}_a^{+}(\omega)$. As before, Equation (20) can be rewritten in terms of the phase and amplitude of the nonlinear response functions,

$$\phi_{\bar{a}}^{+}(\omega) = -\phi_s^{+}(\omega), \tag{21a}$$

$$\alpha_1 N = \frac{1}{|\tilde{R}_s^{+}(\omega)||\tilde{R}_{\bar{a}}^{+}(\omega)|}. \tag{21b}$$

Formally, Eq. (21a) can be used to determine the frequency shift at a finite amplitude of oscillation, while Eq. (21b) relates the oscillation amplitude to the cell density $N$. Since the amplitudes enter quadratically into the nonlinear response functions, they are expected to increase as $(N - N_o)^{1/2}$ just above the threshold cell density $N_o$, e.g., the transition is of the Hopf bifurcation type.

In the Main Text, we have considered the case $c_3 = 1$ and $d_3 = 0$. Numerically, the oscillation frequency is found to decrease as the coupling strength $\bar{N} \equiv \alpha_2\alpha_1 N$ increases (see also Supplementary Figure **1**$A$). This is consistent with Equation (21a) whose solution at selected oscillation amplitudes is shown in Fig. 3$F$ in the Main Text. As the amplitude of the oscillations increase, $\phi_a^{+}(\omega)$ decreases on the low frequency side. Consequently, the intersection point with $\phi_s^{+}(\omega) = \phi_s(\omega)$ shifts to lower frequencies.

Interestingly, the limit cycle associated with the oscillating state in this model shrinks to a fixed point when $\bar{N}$ exceeds an upper threshold value $\bar{N}_b$. The dependence of $\bar{N}_b$ on the adaptation error $\epsilon$, which is assumed to be small, can be estimated as follows. At a fixed point of Eqs. (13) at $d_3 = 0$, we have $s \simeq \alpha_1 N a$ (from $\dot{s} = 0$), $y \simeq \alpha_2 s$ (from $\dot{a} = 0$), and $a \simeq \epsilon y$ (from $\dot{y} = 0$). Consequently, the upper threshold for oscillations has the scaling

$$\bar{N}_b \sim \frac{1}{\epsilon}. \tag{22}$$

Figure **1**$B$ shows the numerical values for $\bar{N}_o$ and $\bar{N}_b$ against the adaptation error $\epsilon$ obtained in our simulations, which confirms (22). The oscillating state expands over a larger range of cell densities when individual cells are more adaptive.

Next, consider the case of nonlinear signal relaxation ($d_3 = 1$) and linear adaptation ($c_3 = 0$). The onset of collective oscillations is similar to the previous case (Supplementary Figure **2**$A$), except that oscillations speed up as the cell density increases further. From Equation (19), we obtain

$$\phi_s^{+}(\omega) = -\arg \tilde{R}_s^{+}(\omega) = -\arctan\Big[\frac{\omega}{\omega_s(1 + 3d_3|\tilde{s}|^2/4)}\Big], \tag{23}$$

which decreases as the oscillation amplitude increases. As shown in Supplementary Figure **2**$C$, the intersection point shifts to the right. The predicted signal oscillation amplitude $B = |\tilde{s}|$ and frequency shift agree quantitatively with our numerical results (Supplementary Figure **2**$C$).

In the more general case when both $c_3$ and $d_3$ are nonzero, we need to first express $\tilde{s}$ and $\tilde{a}_j$ in terms of a common variable that specifies oscillation amplitude before applying the phase-matching condition Equation (21a). As we see from the discussions above, depending on which of the two cubic nonlinearities is stronger, the oscillation frequency may shift either to lower or higher values. In general, nonlinearities may also be present in the dynamics of other intracellular variables which need to be dealt with case by case.

When quadratic nonlinearities are present in the system dynamics, the second harmonic is generated and need to be considered in the perturbative analysis. Consider for example the equation for $a_j$ with an extra term $c_2 a_j^2$. Following the same procedure that led to Eqs. (15), we find an additional term $c_2\tilde{a}_j^*\tilde{a}_j^{(2)}$ on the right-hand side of Equation (15a), where $\tilde{a}_j^{(2)}$ is the amplitude of the second harmonic in $a_j(t)$ (including phase). The amplitude equation for the second harmonic relates $\tilde{a}_j^{(2)}$ to $c_2\tilde{a}_j^2$ and $\alpha_2\tilde{s}^{(2)}$. Together with the equation for $\tilde{s}^{(2)}$, amplitudes of the second harmonic can be expressed as a linear combination of terms $c_2\tilde{a}_j^2$ from different cells. The upshot of this exercise is that coefficient of the cubic term $|\tilde{a}_j|^2\tilde{a}_j$ in Equation (15a) should contain additional contributions proportional to $c_2^2$. The nonlinear response functions (17) and (19) on the limit cycle can still be defined in the same way, and Equation (20) still holds formally. Through $\tilde{s}^{(2)}$, terms $|\tilde{a}_k|^2$ from other cells enter the expression for $\tilde{R}_{a,j}^{+}(\omega)$. Two conclusions can be drawn from this fact: i) as in the case of cubic nonlinearities, the transition is still of the Hopf bifurcation type; ii) $\tilde{R}_{a,j}^{+}(\omega)$ can no longer be determined by simply measuring the response of a given cell to a sinusoidal stimulus at finite strength, as it is affected by the oscillation pattern of other cells in the system due to the quadratic nonlinearity.

#### Supplementary Note 3. Minimal rate of signal decay/clearance at the onset of collective oscillations

Experiments have indicated that sufficiently fast breakdown of the signalling molecule is needed for DQS in dicty[12] and for sustained oscillations in yeast cell suspensions[13]. Below we derive an upper limit for the signal relaxation time

$\tau_s$ to satisfy the phase matching condition Equation (4) in the Main Text. The result is inversely proportional to the adaptation error $\epsilon$ of the intracellular circuit.

Taking Equation (6) for the signal phase shift, $\phi_s = -\tan^{-1}(\omega\tau_s)$, we see that a longer $\tau_s$ yields a larger signal delay $|\phi_s|$ at a given frequency. This is illustrated by Supplementary Figure **3***A*, upper panel, where the horizontal frequency axis is shown on logarithmic scale. For a given $\epsilon$, the intersection of the two phase-shift curves moves to the left, yielding a lower onset oscillation frequency $\omega_o$ and a larger coupling strength $\bar{N}$ (Supplementary Figure **3***B*). When $\tau_s$ reaches beyond an upper limit $\tau_s^*(\epsilon)$, the solution disappears (Supplementary Figure **3***B*). Interestingly, a reduction of $\epsilon$ in Equation (10) increases $\phi_a(\omega)$ on the low frequency side, and rescues the solution (Supplementary Figure **3***A*, lower panel).

The observed inverse dependence of $\tau_s^*$ on $\epsilon$ in Supplementary Figure **3***C* can be justified from the behaviour of the two phase-shift functions at low frequencies. Close to $\omega = 0$, Equation (9) in the Main Text yields $\tilde{R}'_a(\omega) \propto \epsilon$ while $\tilde{R}''_a(\omega) \propto \omega$, as $\tilde{R}''_a$ must be an odd function of $\omega$. This is confirmed by expanding Equation (10) in the Main Text at $\omega = 0$. Consequently, $\phi_a(\omega) \simeq -\tilde{R}''_a/\tilde{R}'_a \propto \omega/\epsilon$. On the other hand, $\phi_s(\omega) \approx -\omega\tau_s$ in this regime. The two curves has an intersection at low frequencies provided

$$\tau_s < \tau_s^* \propto 1/\epsilon. \tag{24}$$

Using the explicit expressions Eqs. (3) and (10) in the Main Text, we obtain $\tau_s^* \simeq \tau_y/\epsilon$ for small $\epsilon$. The critical cell density and onset oscillation frequency at this maximal $\tau_s$ are given approximately by $N_o \simeq K(\alpha_1\alpha_2\epsilon)^{-1}$ and $\omega_o \simeq \epsilon\tau_y^{-1}$, respectively. This is compared to $N_o \simeq K(\alpha_1\alpha_2)^{-1}$ and $\omega_o \simeq (\tau_a\tau_y)^{-1/2}$ at $\tau_s < (\tau_a\tau_y)^{1/2}$, which are insensitive to $\epsilon$ as long as it is sufficiently small.

#### Supplementary Note 4. Yeast glycolysis: single-cell perturbation study in the full model

The full intracellular reaction network of the kinetic model by du Preez et al.[15] is shown in Supplementary Figure **4**. It contains around 20 reactions and 15 metabolite concentrations as dynamical variables. The reaction fluxes are highly nonlinear functions of these variables. Predictions of the model were shown to agree semi-quantitatively with experimental data on yeast glycolytic oscillations[16]. Below we use the same parameter values as adopted in the original model termed dupree2 in[15], unless otherwise stated.

Supplementary Figure **5** shows an oscillatory solution of the model at the glucose concentration Glco =10 mM. The oscillation frequency is $\omega_0 \approx 21$ min$^{-1}$, corresponding to a period of $\tau_0 \simeq 0.3$ min. The mean concentration of ACE is 0.17 mM (Supplementary Figure **5***B*). Reaction fluxes are concentrated along the linear pathway from Glco to ETOH, while the side reactions carry much smaller flux (Supplementary Figure **5***C*, blue box). Below, we present response properties of the model using ACE concentration as the second control variable, in addition to the extracellular glucose concentration. Time is measured in minutes and concentrations in mM.

To move out of the oscillatory regime, we lower the mean acetaldehyde concentration to $\mathrm{ACE}_0 = 0.05$. Experimentally, this can be achieved by adding cyanide (KCN) which reacts with ACE in the solution[17]. Supplementary Figure **6** shows the time course of metabolites under a step-wise increase in the ACE concentration. The four panels are organised following the order of metabolites along the glycolytic pathway, with the addition of ATP, ADP and NAD. Most metabolites adapt at least partially, except F16P and TRIO upstream of the reaction GAPDH that uses NAD and NADH as cofactors. The redox pair NAD and NADH, being tightly connected to ACE, do not adapt either.

We now consider oscillations of the same set of metabolites stimulated by a periodic redox signal at the frequency $\omega_0$ of spontaneous oscillations mentioned above. In Supplementary Figure **7***A*, ATP, G6P and F6P are approximately in phase with each other, but they are out of phase with Glci at the entry point of the pathway. The non-adaptive F16P has a behaviour of its own. The phase relations for these metabolites have been measured experimentally, and the results agree well with our numerics[18]. In Figs. **7***B*-*C*, metabolites from BPG down to PEP share nearly the same phase with each other and with ATP. The non-adaptive TRIO lags slightly behind F16P. In Supplementary Figure **7***D*, PYR at the end of the glycolytic pathway has an approximately 90° phase lead over ATP, and furthermore a smaller phase lead over ACE and NAD.

Supplementary Figure **8** shows the phase shifts of metabolites against a sinusoidal signal ACE obtained from our numerical simulations over a broad frequency range. Apart from PYR, the phase relationships among metabolites at $\omega_0$ hold also at lower frequencies. In Supplementary Figure **8***B*, it is seen that NAD has essentially the same phase as ACE in the frequency interval, while NADH is completely out of phase. Therefore, on the timescale $\tau_0$, the phase information of ACE is passed without delay onto the redox ratio NAD/NADH, and fed into the network through the reaction GAPDH. Around $\omega_0$, the phase lead of NADH over ACE is slightly below 180°, as observed in experiments on glycolytic oscillations[19]. Supplementary Figure **8***C* shows the downstream metabolites from BPG to PEP oscillate in phase with each other for $\omega \leq \omega_0$, meaning the internal time scales for this part of the pathway are shorter than $\tau_0$. In contrast, PYR develops a phase lead in the intermediate frequency regime, as indicated by the two black arrows in Supplementary Figure **8**$C$. (Note that in Fig. 5$D$ of the Main Text, the phase lead extends to zero frequency indicating that the width of the regime depends on the glycolytic flux.) The adaptive variable ATP also has a phase lead in the entire low frequency region.

Supplementary Figure **8**$D$ shows the following phase relations between ATP and several other metabolites as summarized by the equations below,

$$\phi_{\mathrm{ATP}} = \pi + \phi_{\mathrm{ADP}} = \pi + \phi_{\mathrm{AMP}} \approx \phi_{\mathrm{BPG}} \approx \phi_{\mathrm{PEP}} \approx \pi + \phi_{\mathrm{GLCi}}. \tag{25}$$

The first two relations among the nucleotides ATP, ADP and AMP simply reflect the conservation of their total number, and that the fraction of AMP is much lower than the other two. In-phase relations apply to substrates BPG and PEP of the ATP harvesting reactions PGK and PYK, respectively, while the out-of-phase relation is observed for GLCi in the ATP consuming reaction GLK. The fact that these relations hold almost strictly in the entire frequency region suggests that quasi-steady-state conditions apply to these and neighbouring reactions. It also suggests a prominent role of ATP in synchronising the phase of metabolites distributed along the glycolytic pathway.

In summary, our numerical results suggest the following mechanism of adaptation. Under a stepwise increase of ACE concentration, the information is passed with negligible delay to the redox ratio NAD/NADH, and then through the delayed reaction GAPDH to BPG and PEP, transiently boosting ATP production. The transient increase of ATP concentration then reduces the upstream glycolytic flux by inhibiting the reaction PFK, which in turn decreases the downstream TRIO concentration, eventually returning the GAPDH flux to its pre-stimulus level. Although many metabolites adapt, the negative feedback loop of ATP production appears to be the core. Fig. 5$B$ in the Main Text shows a more complete phase diagram of the response properties at other values of $\mathrm{ACE}_0$ and Glco concentrations, including the region of spontaneous oscillations.

### Supplementary Note 5. A minimal model for ATP autocatalysis coupled to intracellular redox state

We constructed a minimal model to test various quantitative aspects of the adaptation mechanism described above. Reduction in the number of dynamic variables is achieved by lumping consecutive metabolites along the linear pathway that are phase synchronised into a single variable denoting their total concentration. This is a reasonable approximation when interconversion among these metabolites is much faster than the time of interest, e.g. the oscillation period. Supplementary Figure **9**$A$ illustrates the selected variables and their interactions. Here, $y$ represents intermediate metabolites that do not adapt (F16P and TRIO), thereby playing the role of a memory node. The variable $z$ represents metabolites from BPG to PEP along the glycolytic pathway. The ATP concentration is denoted by $p$, while the concentration of PYR, substrate for the ACE producing reaction PDC and thus the corresponding cell activity here, is denoted by $a$. Since NAD (NADH) is always in phase (out of phase) with ACE, we adopt the redox ratio NAD/NADH as the signal $s$ instead. Motivated by a phenomenological two-component model for glycolytic oscillations in Ref.[20], we introduce a minimal model of glycolysis with redox control as follows:

$$\tau \dot{y} = \frac{2p}{1+p^{2h}} - (\alpha_2 s + c_0) y - \epsilon y, \tag{26a}$$

$$\tau \dot{z} = (\alpha_2 s + c_0) y - \frac{2z}{1+p^2}, \tag{26b}$$

$$\tau \dot{p} = -\frac{2p}{1+p^{2h}} + 2\frac{2z}{1+p^2} - \frac{2p^2}{1+p^2}. \tag{26c}$$

Here, $2p/(1+p^{2h})$ gives the reaction flux of PFK that consumes ATP and is also inhibited by ATP at high concentrations (i.e., the negative feedback loop), with the inhibition strength set by the exponent $h(> 1/2)$. The entry carbon flux into the glycolysis pathway is assumed not to be rate limiting, e.g.., one is in a situation of high extracellular glucose concentration. The term $(\alpha_2 s + c_0)y$ gives the reaction flux of GAPDH, where $c_0$ sets the "basal" enzyme velocity at $s = 0$. Leakage of TRIO into the side branch is represented by $\epsilon y$. The term $2z/(1+p^2)$ gives the reaction flux of PYK (and also PGK), which produces ATP but is also inhibited by ATP. In Equation (26c), the stoichiometric factors 1 and 2 in the first two terms on the right-hand-side correspond to the ATP consumption and production upstream and downstream of TRIO, respectively. ATP consumption by the cell outside of glycolysis (e.g., ATPase activity) is modelled by the term $2p^2/(1+p^2)$, which grows with the ATP concentration until saturation at a maximal value 2. The output variable $a$ (PYR) is produced by the same flux that produces $p$ (ATP) and degraded at a constant rate $\alpha_1$,

$$\tau \dot{a} = \frac{2z}{1+p^2} - \alpha_1 a. \tag{26d}$$

The glyoxylate shunt GLYO, which is active at $\mathrm{ACE}_0 \simeq 0.2$ mM or above in the full model, is turned off. For simplicity, we have chosen the time constants on the left-hand-side of the equations to be the same. As we show below, this choice is adequate for recovering the main low frequency properties of the full model.

Supplementary Figure **9***B* shows the response of the dynamical variables to a sinusoidal redox variation centred around $s_0 = 0.04$. Except the buffer variable $y$, all other variables show adaptive behaviour, with $a$ gaining a phase lead of 90° over $p$. For comparison, we show in Supplementary Figure **9***C* the response properties of corresponding metabolites in the full model in the adaptive regime, which are indeed quite similar. We have examined the response properties of the minimal model at other values of $s_0$ and identified four qualitatively different regimes as shown in Supplementary Figure **9***D*. As in the case of the full model with sufficient glucose (Main Text, Fig. 5*B*), spontaneous oscillations (i.e., limit cycle solution) occur at intermediate values of $s_0$, flanked by adaptive but non-oscillatory regions.

We note in passing that the two-component model of Chandra *et al.*[20] also exhibits spontaneous oscillations when the rate constant $k$ of the pyruvate kinase reaction (PYK in Supplementary Figure **9***A*) takes on intermediate values. As $k$ affects the delay time of the negative feedback control in ATP production, in this sense it plays a similar role as $s_0$. However, our model contains an additional buffer node TRIO which is necessary for the adaptive behaviour seen in Supplementary Figure **9**. We have also made the ATP consumption rate dependent on the ATP concentration to eliminate certain pathological aspects of the Chandra *et al.* model at low values of $p$. Furthermore, our numerical analysis suggests that a sufficiently small but finite adaptation error $\epsilon$ associated with low flux diversion is needed to reproduce the response diagram Supplementary Figure **9***D*. On the high (oxidative) end of $s_0$, the reaction GAPDH drives down $y$ (TRIO) and hence the flux of the side reaction, making the system adaptive even when $\epsilon \sim 1$.

Comparing the response diagrams of the minimal model (Supplementary Figure **9***D*) and of the full model at high extracellular glucose concentrations (Main Text, Fig. 5*B*), we see that the adaptive regime on the oxidative side is restricted to a much narrower region in the latter case. Upon a detailed investigation of the full model we found that, at higher values of $\mathrm{ACE}_0$, the side reaction GLYO is activated. Shutting down the reaction, we obtained a response diagram similar to that of the minimal model (Supplementary Figure **10**). The reaction GLYO uses NAD as cofactor and consumes ATP (see Supplementary Figure **4**). With regard to the change in NAD/NADH ratio upon an upshift of ACE, it has an opposite effect as compared to ADH. This and ATP consumption by GLYO leads to a sign reversal in the transient response of ATP to ACE upshift at $\mathrm{ACE}_0 \simeq 0.2$ mM in the full model (Fig. 5*B* in the Main Text). Inhibition of GLYO eliminates the sign switch and makes PYR adapting to ACE over a much larger region of the phase diagram (Supplementary Figure **10**).

Supplementary Figure **11** shows representative time courses of the PYK reaction flux to a stepwise ACE signal, computed using the original and modified glycolysis model, as well as the minimal model. Concentration of its product, PYR, is found to be proportional to the PYK reaction flux in all three models, i.e., the degradation rate of PYR is a constant. The original and modified models exhibit nearly identical adaptive response on the low ACE (reductive) side, but differ on the high ACE (oxidative) side. In the latter case, the PYK flux is significantly higher and also non-adaptive when the glyoxylate shunt (GLYO) is on. Further numerical investigations of the full model with blocked GLYO reaction show that it shares the following features of the minimal model as the oxidation level increases: 1) the frequency inside the oscillatory regime increases; 2) (mean) $p$ (ATP) and $z$ (BPG, P3G, P2G, and PEP concentrations) increase by a moderate amount; 3) $y$ (TRIO and F16P concentrations) decreases; 4) $a$ (PYR concentration) first increases, then decreases. Experimental time-course measurement with blocked glyoxylate shunt will serve to validate or improve the model assumptions.

#### Supplementary Note 6. A model for glycolytic oscillations in yeast cell suspensions

To study collective oscillations in a population of cells whose internal dynamics follows Eqs. (26), we adopt the following signal dynamics as in Ref.[21]:

$$\tau_s \dot{s}_{\mathrm{in}} = \alpha_1 a - k_{\mathrm{in}} s_{\mathrm{in}} - D(s_{\mathrm{in}} - s_{\mathrm{ex}}), \tag{27a}$$

$$\tau_s \dot{s}_{\mathrm{ex}} = \rho D(s_{\mathrm{in}} - s_{\mathrm{ex}}) - k_{\mathrm{ex}} s_{\mathrm{ex}}. \tag{27b}$$

Here $s_{\mathrm{in}}$ and $s_{\mathrm{ex}}$ are the intracellular and extracellular signal concentration, respectively; $D$ is the membrane permeability of the signalling molecule; $k_{\mathrm{in}}$ and $k_{\mathrm{ex}}$ are the intracellular and extracellular signal degradation rate; and $\rho$ is the volume fraction of yeast cells, which increases with the cell density, and saturates at 1. Hereafter, we call $\rho$ the cell density directly. In the steady-state, the extracellular signal strength (i.e., acetaldehyde concentration) is a function of $\rho$ and $D$.

Let us first consider the situation of fast equilibrium between $s_{\mathrm{in}}$ and $s_{\mathrm{out}}$. Previously, Silvia De Monte *et al.* proposed a diffusion timescale $\tau_s \approx 0.003$ s by assuming a quasi-stationary concentration profile and that ACE molecules need to diffuse across a spherical shell with an inner radius $r_1 = 3$ $\mu$m and an outer radius $r_2 = 6.5$ $\mu$m[22].

This diffusion timescale is much smaller than the oscillation period of 37 s. Assuming the time for an ACE molecule to cross the cell membrane is of the order of 1 s or less, we obtain the following approximate equation for $s = (s_{\rm in}+s_{\rm ex})/2$,

$$\tau_s \dot{s} = \frac{\rho}{1+\rho}\alpha_1 a - \Big(\frac{\rho k_{\rm in} + k_{\rm ex}}{\rho+1}\Big)s. \tag{28}$$

Up to corrections of order $\epsilon$, the stationary state of the dynamical system defined by Eqs. (26) and (28) is given approximately by,

$$p \approx 1, \quad z \approx 1, \quad y \approx \frac{1}{\alpha_2 s + c_0}, \quad a \approx \frac{1}{\alpha_1}, \quad s \approx \frac{\rho}{\rho k_{\rm in} + k_{\rm ex}}. \tag{29}$$

The signal strength increases with the cell density and saturates at $1/(k_{\rm in} + k_{\rm ex})$. At small but finite $\epsilon$, corrections to the above expressions become significant at large $y$ or small $s$, where the side reaction G3PDH in Supplementary Figure **4** is activated to divert the glycolytic flux. In the numerical studies presented below, we set the two ACE degradation rates $k_{\rm in}$ and $k_{\rm ex}$ to be small. The signal strength $s$ varies over a broad range as the cell density $\rho$ increases.

Supplementary Figure **12** shows numerical solutions of the coupled minimal model at four selected $\rho$ values. Except the case at $\rho = 0.01$, oscillations of $s$ and the intracellular variables are seen. In Supplementary Figure **13**, we plot the oscillation amplitudes and time-averaged values of $s$ and $a$ against the cell density $\rho$. From the lower panel of Supplementary Figure **13**$A$ we see that, for the signal dynamics chosen, the lower adaptive regime in Supplementary Figure **9**$D$ is mapped to a narrow interval of cell density $0.003 < \rho < 0.013$. From Supplementary Figure **12**, we see that onset of collective oscillations in the coupled system takes place somewhere between $\rho = 0.01$ and 0.014. More detailed studies indicate that the transition is not the expected Hopf bifurcation type, but instead emergence of a limit cycle at finite amplitude. Similar behaviour was seen in the study of the full kinetic model (see Fig. 10 in[16]). On the other hand, experimental work seem to support the Hopf bifurcation scenario[22,23]. The discrepancy could be due to cell-to-cell variability in the experimental system.

Beyond the onset point, oscillation amplitudes vary continuously with the cell density. For $\rho > \rho_c = 0.34$, the time-averaged value of $s$ falls in the upper adaptive regime in Supplementary Figure **9**$E$. Since the cell density here already exceeds the threshold value required for collective behaviour of adaptive units, oscillations continue.

Finally, we present numerical results demonstrating the effect of a slower cross-membrane transport of acetaldehyde on the collective dynamics. The system dynamics is defined by Eqs. (26) for individual cells (with $s = s_{\rm in}$) together with Eqs. (27) for the intracellular and extracellular signal concentrations. Supplementary Figure **14** shows the oscillation amplitude of $s_{\rm in}$ together with the time-averaged values of $s_{\rm in}$ and $s_{\rm out}$ at selected values of $D$. At $D = 100$, $s_{\rm in}$ and $s_{\rm out}$ are nearly identical and the system behaviour is essentially the same as described above under the fast equilibrium assumption. At $D = 1$, the time-averaged value of $s_{ext}$ is noticeably smaller than that of $s_{\rm in}$, indicating a significant gradient of acetaldehyde concentration across the cell membrane. Nevertheless, collective oscillations via DQS continue at all cell densities. As we further reduce $D$, DQS stops at large cell densities, as illustrated by the case for $D = 0.4$. Collective oscillations disappear at $D = 0.1$. Here, $s_{\rm in}$ remains high due to the slow intracellular degradation rate $k_{\rm in}$, which places the single-cell dynamics in the upper adaptive regime even when the cell is isolated. However, the phase delay across the cell membrane changes the response properties of the cell to external signal variations. In this case, $s_{\rm in}$ should be considered as the sender of the external signal but as one can see from Eq. (27a), the adaptation (phase lead) of $a$ to $s_{\rm in}$ does not translate to phase lead of $s_{\rm in}$ to $s_{\rm ex}$ when $D$ is small. The latter is required for the adaptation route to collective oscillations.

Now, we apply our linear response theory to predict the disappearance of DQS in this system at $D = 0.4$, where the oscillation amplitude decreases continuously to zero at large cell densities (Supplementary Figure **14**). The onset condition cannot be predicted from this theory as collective oscillations emerge first through synchronisation of individual oscillators. Mapping this model to our generic framework, the intracellular signal $s_{\rm in}$ is the "activity", while the extracellular signal $s_{\rm ex}$ is the mediating signal. Absorbing the contribution of cell density into the signal response, i.e.,

$$\tilde{R}_{s_{\rm ex}}(\rho,\omega) \equiv \frac{\tilde{s}_{\rm ex}}{\tilde{s}_{\rm in}} = \frac{\rho D}{k_{\rm ex} - i\omega\tau_s + \rho D}, \tag{30}$$

as computed from Equation (27b) using Fourier transform, the self-consistency for collective oscillations in this example requires

$$\phi_{s_{\rm in}}(\rho_c,\omega_c) = -\phi_{s_{\rm ex}}(\rho_c,\omega_c), \qquad |\tilde{R}_{s_{\rm in}}(\rho_c,\omega_c)\tilde{R}_{s_{\rm ex}}(\rho_c,\omega_c)| = 1. \tag{31}$$

A simple form of signal relay efficiency no longer exists due to the nonlinear effect of cell density on the signal spectrum. Here, $\tilde{R}_{s_{\rm in}} \equiv \tilde{s}_{\rm in}/\tilde{s}_{\rm ex}$ measures how an individual cell, defined by Equation (26) and Equation (27a), responds to a periodic perturbation of $s_{\text{ex}}$ at frequency $\omega$. As glycolysis is a nonlinear process, $\tilde{R}_{s_{\text{in}}}$ also depends on the average level $\bar{s}_{\text{ex}}$ of extracellular signal at which small periodic stimulation is superimposed. As we are more interested in the response spectrum at its natural context of interacting populations, we perturb the cell at $\bar{s}_{\text{ex}}$ determined from the population dynamics at a given cell density $\rho$. Hence, the signal-level dependence of $\tilde{R}_{s_{\text{in}}}$ translates into cell-density dependence. Given the numerically computed two-dimensional spectrum $\tilde{R}_{s_{\text{in}}}(\rho,\omega)$ and $\tilde{R}_{s_{\text{ex}}}(\rho,\omega)$, the critical (fixed) point $\rho_c$ can be found via numerical iteration. At the critical cell density $\rho_c$, we expect the two equations in Equation (31) to be satisfied at the same frequency. This is indeed observed in Supplementary Figure **15***A*. Note that the phase shift $\phi_{s_{\text{in}}}$ is positive only in the intermediate frequency regime, unlike the case for typical adaptive variables seen in other examples in this paper. While $\phi_{s_{\text{in}}}$ has a phase-leading regime induced by the upper-stream adaptive variable $a$, $s_{\text{in}}$ may not necessarily be adaptive to $s_{\text{ex}}$, especially at large $D$, where $s_{\text{in}}$ and $s_{\text{ex}}$ would be almost identical at any time. The predicted critical cell density and frequency agree well with results from direct simulation of the population dynamics (Supplementary Figure **15***B*).

In summary, under fast equilibration between intracellular and extracellular acetaldehyde concentrations, the coupled system exhibits collective oscillations over a broad range of cell densities, encompassing the adaptive and oscillatory regimes of a single cell. Onset of collective oscillations at low cell densities exhibit complex behaviour due to the assumed sensitivity of the reaction GAPDH to the NAD/NADH ratio. Delay in the cross-membrane transport of acetaldehyde weakens adaptation of intracellular metabolite concentrations to change in the extracellular acetaldehyde concentration, and may eliminate collective oscillations altogether when the delay is too long[13]. At moderate delays, rise in the intracellular acetaldehyde concentration brings individual cells to the oscillatory regime even when in isolation. The enhanced oscillation amplitude at $D = 1$ and low cell densities seen in Supplementary Figure **14**, however, is obtained under the assumption that all cells in the population behave identically. This behaviour is susceptible to cell-to-cell variations as well as temporal noise in intracellular dynamics. As $D$ decreases further, an isolated cell exits from the oscillatory state and moves into the upper adaptive region, as shown in the last two columns in Supplementary Figure **14**. Collective oscillations at $D = 0.4$ is attributed entirely to DQS. Our model study exposes this and other subtleties that can affect interpretation of collective oscillations. The specific effects we identified in this work could serve to guide the design of future experiments where various model parameters can be controlled quantitatively, e.g., $k_{\text{ex}}$ for extracellular degradation rate of acetaldehyde by adjusting the flow rate in microfluidic setups[17].
